# Directed evolution and characterisation of light harvesting complexes with altered energy transfer dynamics in purple non-sulfur bacteria

**DOI:** 10.1101/2025.10.01.678743

**Authors:** Eveliny Tomás Nery, Claudio Avignone-Rossa, Luca Sapienza, Johnjoe McFadden, Alexandra Olaya-Castro, José I. Jiménez

**Affiliations:** The Leverhulme Quantum Biology Doctoral Training Centre, University of Surrey, GU2 7XH, Guildford, UK; Department of Neuroscience and Biomedical Engineering, Aalto University, 02150, Espoo, Finland; School of Biosciences, Faculty of Health and Medical Sciences, University of Surrey, GU2 7XH, Guildford, UK; Department of Engineering, University of Cambridge, Cambridge, CB3 0FA, United Kingdom; Department of Physics and Astronomy, University College London, Gower Street WC1E 6BT, London, UK; Department of Life Sciences, Imperial College London, SW7 2AZ, London, UK; Imperial Centre for Engineering Biology, Imperial College London, SW7 2AZ, London, UK

**Keywords:** Photosynthesis, quantum coherence, purple non-sulfur bacteria (PNSB), light-harvesting, *Rhodobacter sphaeroides*, LH2

## Abstract

Purple non-sulfur bacteria (PNSB) are metabolically versatile microorganisms that inhabit diverse environments by taking advantage of a remarkably efficient photosynthetic machinery. In this work, we describe the creation of a workflow for experimental studies that aim to compare the energy transfer mechanisms occurring within structural variants of the light-harvesting complex 2 (LH2) of PNSB. Through the creation of a library of LH2 variants using site-directed mutagenesis, we engineered proteins with different spectral properties to be expressed in a LH2-defficient mutant of the model PNSB *Rhodobacter sphaeroides*. We validated this approach by reproducing a previously described mutant exhibitng a blue-shift, in addition to identifying a novel mutant exhibiting a red-shift of the B850 absorption peak. We characterised the fluorescence lifetime of the purified LH2 spectral variants *in vitro*, and performed a bacterial growth assay to assess the fitness of the LH2 variants *in vivo* under oversaturating light conditions. Our results suggest that the LH2s variants expressed by PNSB in nature reflect the intricate tunning of their quantum properties not towards the fastest energy transfer but towards the optimum light-harvesting efficiency which is defined by diverse environmental factors.

**Statement of significance:** In this work we report a platform for the systematic investigation of the mutational landscape of light harvesting complexes forming part of the photosynthetic machinery of purple non-sulfur bacteria. By conducting directed evolution of selected residues of the light-harvesting antenna (LH2) of *Rhodobacter sphaeroides* we identified a novel mutant with distinct and red-shifted spectral properties. We characterised the energy transfer dynamics of this and a previous characterised mutant and demonstrated that the new variant confers a phenotypic growth advantage when cultured with high-intensity light. Our findings offer new insights into the mechanisms of light capture and energy transfer also bridging the *in vitro* observations with quantifiable fitness advantages under the conditions tested.

## Introduction

Purple non-sulfur bacteria (PNSB) is a group of metabolically versatile photosynthetic alpha- and beta-proteobacteria which inhabits a vast range of aquatic and terrestrial environments (1). In addition to their importance as primary producers and their potential in the development of biotechnological products (2, 3), this group of bacteria has gained special attention in the last decades due to the unique characteristics of the energy transfer dynamics happening within their light-harvesting complexes (LHCs; (4–6)). Most PNSB perform photosynthesis when grown under anaerobic conditions and express two types of LHCs: LH1, which contains the reaction centre (RC) – referred to as LH1-RC – and LH2, responsible for absorbing photons and channelling electronic excitation energy towards the LH1-RC (7, 8). Both LHCs consist of heterodimers of the non-covalently bound subunits α (alpha) and β (beta), which associate with bacteriochlorophylls (bchls) and carotenoids (cars; (7, 9)).

The subunits forming the LH2 of PNSB, also called apoproteins, are encoded by the *pucBA* genes (8). This protein features two rings of bchls which exhibit distinct spectral properties: B800, responsible for an 800 nm absorption band; B850, with a strong excitonic coupling between pigments that results in an absorption peak near 850 nm (Fig. 1; (10)). Some PNSB species modulate the spectral properties of their LH2 when grown under different light conditions by expressing variants of the *pucBA* genes, such as those encoding the low-light variant in *Rhodoblastus acidophilus*, also called LH3, which displays a 30 nm blue-shift in its B850 absorption peak (11, 12). These shifts arise from changes in site energies, pigment-pigment interactions, and charge-transfer (CT) states (13–15). However, the physiological relevance of these shifts for PNSB and their evolutionary significance remain speculative, since most studies on LH2 function rely on *in vitro* models or computational simulations (16– 18). In this work we develop a platform that addresses this knowledge gap by enabling the recombinant expression of semi-randomly engineered LH2 variants in the model PNSB *Rhodobacter sphaeroides* (*Cereibacter sphaeroides*; (19)).

**Figure 1.**
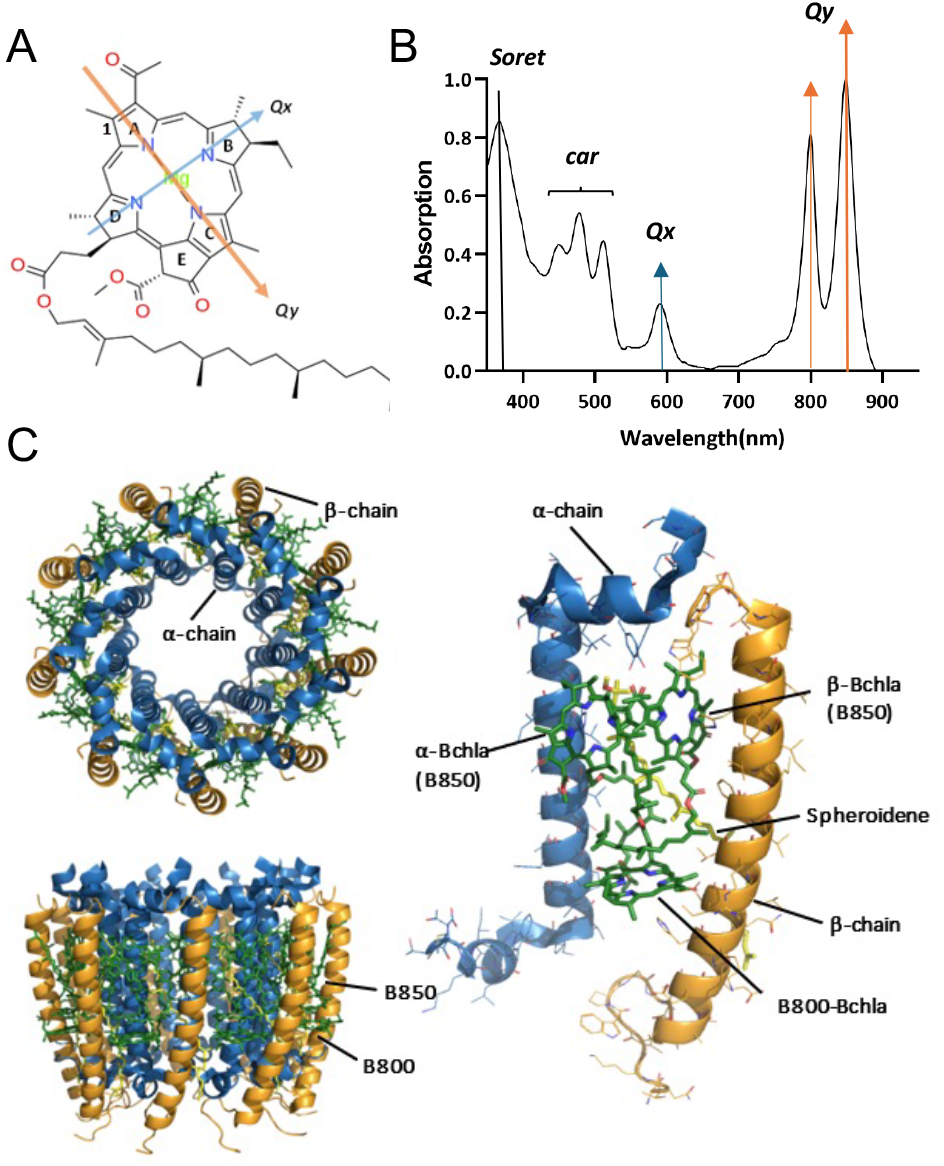
Error! No text of specified style in document. Structure and absorption spectrum of the LH2 of *Rb sphaeroides*. **A**. Mg^2+^ coordination in bchlas showing the interactions responsible for the *Qx* and *Qy* absorption bands (diagram extracted from Wikipedia, public domain). **B**. Absorption spectrum of native LH2 complexes purified from *Rb. sphaeroides*. The peaks resulting from the carotenoids and the Soret and Q bands of the bchla constituting this protein are indicated. **C**. Cryo-Electron Microscopy (EM) structure of *Rb. sphaeroides* 2.4.1 LH2 complex (PDB entry: 7PBW). Left: Top (top; periplasmic side) and side (bottom) views of the protein displaying the α and β apoproteins in blue and orange, respectively, in addition to the bchls in green and the carotenoids in yellow. The positions of the B850 and B800 rings of bchlas are indicated. Right: LH2 subunit composed by a pair of apoproteins α and β. In this figure the bchls composing the B850 (α and β-Bchla) and B800 (B800-Bchla) rings and the carotenoid spheroidene, which is produced by *Rb. sphaeroides*, are indicated. The bchls’ central Mg^2+^ have been omitted for clarity. The structure representation was created with Pymol.

A reason for focusing on *Rb. sphaeroides* is that the structure of their LH2 complexes is very well known. It comprises 9 αβ heterodimers binding 27 bchlas and 9 spheroidene cars (Fig. 1; (20)). The B800 pigments are held in place by hydrogen bonds extending from their central Mg^2+^ (Fig. 1A), and their spectral properties (Fig. 1B) and pigment specificity can be modulated by modifications in nearby residues such as β-His22 and β-Arg30 (21, 22). The B850 ring, composed of 18 bchlas, features strong excitonic coupling and superradiant properties (10, 13, 20, 23). The coordination of these molecules within the protein scaffold is obtained through hydrogen bonds formed between their central Mg^2+^ and α-His31 and β-His40 residues and between an C3-acetyl group of the α-bchla to the α-Tyr45 and the α-Tyr44 the adjacent heterodimer (Fig. 1C; (20)).

In addition, the existence of LH2-free genetic backgrounds of *Rb. sphaeroides* and the possibility of tuning the spectral properties of the LHCs of PNSB through modifications of their protein scaffold, has allowed studies of their structure-function relationships. For example, the replacement of a histidine at position β-21 with serine causes a 4 nm red shift in the B850 Qy absorption band while substituting β-Arg29 with glutamate results in a significant blue shift and broadening of the B800 Qy band (24). Similarly, substitutions at β-30 lead to spectral shifts in the B800 Qy band, primarily due to changes in site energies from hydrogen bonding between the C3-acetyl group of B800 and the protein scaffold (25).

The occurrence of spectral variants in nature suggests that the LH2 system has been tuned throughout its evolutionary history, primarily through modifications in the composition of its apoproteins. Despite the abundant structural evidence, the *in vivo* implications of spectral shifts are largely missing. Here we present an experimental workflow to design, express and characterise libraries of semi-randomly generated LH2 proteins spanning a wider range of variations than those studied so far. This strategy combines *in vivo* and *in vitro* assays involving genetic engineering, spectroscopic analysis, and bacterial growth studies: this allows exploring how the coupling between pigment and proteins can be modified to generate novel absorption characteristics that impact LH2 energy transfer dynamics and the fitness of bacterial strains expressing such variants under specific light conditions.

Our results show that the random evolution of selected regions in the protein can lead to distinct absorption properties, accompanied by corresponding shifts in emission spectra and fluorescence lifetimes of the antenna protein *in vitro*. These results also suggest a significant impact *in vivo*, particularly on the fitness of bacterial strains expressing these variants when grown under high-light intensities.

## Materials and Methods

### Strains, plasmids and bacterial growth

The wild-type strain of *Rb. sphaeroides* was obtained from the DSMZ culture collection (DSM158). For expression of LH2 variants we used a *Rb. sphaeroides* LH2 deletion mutant (Δpuc1BA; (26)), called Rb.X1 throughout this work. Both strains were cultivated in M22+ medium (27) at 30 °C aerobically in the dark or anaerobically (10% CO2, 10% H2, and 80% N2) in the presence of light (Fig. S1). During cloning procedures, the strains were incubated in aerobic conditions in a rotary shaker at 150 rpm in the dark. For growth and selection assays, the strains were grown statically. M22+ medium was prepared by dissolving KH_2_PO_4_(3.06 g/L), K_4_HPO_4_(3 g/L), DL-lactic acid (2.5 g/L), (NH_4_)_4_SO_4_(0.5 g/L), NaCl (0.5 g/L), Na_4_succinate (4.34 g/L), L-glutamic acid (0.27 g/L), DL-aspartic acid (0.04 g/L), and casamino acids (1 g/L) in distilled water, pH 7.0. For solid media, agar was added at a final concentration of 1.5% (w/v). After autoclaving, the medium was supplemented with 0.1 ml/L of filter sterilized vitamins solution (nicotinic acid 0.1 g/L, thiamine 0.05 g/L, 4-aminobenzoic acid 0.01 g/L, biotin 1 mg/L) and 20 ml/L of Solution C (nitrilotriacetic acid 10 g/L, MgCl_2_24 g/L, CaCl_2_3.34 g/L, EDTA 0.125 g/L, ZnCl_2_0.26 g/L, FeCl_2_0.25 g/L, MnCl_2_0.09 g/L, (NH_4_)_4_Mo_4_O_4_·4H_2_O 9 mg/L, CuCl_4_8 mg/L, Co(NO_4_)_4_1.2 mg/L, orthoboric acid 6 mg/L). *Escherichia coli* strain DH5α was cultured in Lysogeny Broth (LB; Sigma-Aldrich) at 37°C with shaking (150 rpm). When required, kanamycin (km) was added to culture medium at the final concentration of 20 and 50 μg/ml for *Rb. sphaeroides* and *E. coli*, respectively. The plasmid pSEVA221 was used for cloning and recombinant expression in *Rb. sphaeroides* (28).

### Molecular techniques

The native *pucBA* genes with their promoter regions were amplified from the wild-type *Rb. sphaeroides* strain using colony PCR with primers pucBAC_F and pucBAC_R (Table S1). These primers were designed based on the genome sequence of the strain available in NCBI (GenBank: X68796) and included EcoRI and BamHI restriction sites flanking the amplified region. The PCR product was purified using the QIAquick PCR Purification Kit (QIAGEN). The PCR product and pSEVA221 were then digested with the respective endonucleases (EcoRI and BamHI-HF® Restriction Endonucleases, New England Biolabs) and ligated with T4 DNA ligase (New England Biolabs) following the protocol for restriction and ligation provided by the supplier. After ligation, the resulting vector p221BAC was transformed into chemically competent *E. coli* DH5α cells. These cells were then incubated overnight at 37 °C in LB agar plates supplemented with km. Presence of the insert in the resulting colonies was checked by colony PCR using the same pair of primers as above. A positive colony was inoculated into LB supplemented with km plasmids were purified using QIAprep Spin Miniprep Kit (QIAGEN). The presence of the insert in p221BAC was confirmed by Sanger sequencing (GENEWIZ® from Azenta Life Sciences) using the primers F24/R24 (Table S1).

The construct p221BAC was delivered to Rb.X1 by electroporation. In brief, the strain was inoculated in M22+ broth and incubated aerobically in a rotary shaker (150 rpm) at 30 °C until reaching the late-exponential growth phase. Next, approximately 5 mL of culture was centrifuged, resuspended in 1mL of M22+ broth and cooled in ice for 15 min. After cooling, the cells were centrifuged again and washed with 1 ml of cold sterile milli-Q water three times to remove traces of the medium. Then, the cells were resuspended in 100 µL of cold milli-Q water containing ∼1 µg of plasmid and placed into an electroporation cuvette (1 mm gap; Fisherbrand). The electroporation was carried using a Gene Pulser XcellTM electroporator (Bio-Rad Laboratories) configured to deliver a pulse with field strength of 20 kV/cm (2000 V, 200 Ω, 25 µF). Immediately after the pulse, 1 mL of M22+ (at 30°C) was added to the cuvette and the cells were allowed to recover for 12 h aerobically at 30 °C. After that, the cells were spread in M22+ agar supplemented with km and incubated under the same conditions until colonies could be observed.

### Design of LH2 variants

The selection of residues for mutation was informed by the alignment of α and β LH2 subunits expressed by different strains of PNSB using the MEGA software (Fig. S2; (29)) and the Cryo-EM structure of LH2 from *Rb. sphaeroides* (20) (RCSB Protein Data Bank ID 7PBW). Additionally, target amino acids were selected based on their proximity to the porphyrin rings of the bchlas and their likelihood of causing spectral shifts in LH2. After selection, PCR site-directed random mutagenesis of three amino acids of the LH2 apoproteins was carried out by designing pairs of oligonucleotides with 20 bp of homology (Table S1) bearing a random codon (NNN) in the target site at the non-homologous “tail” annealing to the backbone of pSEVA221. The PCR products were obtained using p221BAC as template and then treated with 20 U of the restriction enzyme DpnI (New England Biolabs) per µg of DNA, overnight at 37 °C. These single mutant amino acid libraries were transformed into chemically competent *E. coli* DH5α cells which were then incubated overnight. Colonies were collected from the plates by washing with phosphate buffer saline (PBS) and a library of plasmids was obtained after purification using the QIAprep Spin Miniprep Kit (Qiagen).

The plasmid libraries obtained were transformed into Rb.X1 by electroporation as previously described. The strain was then incubated in anaerobic conditions under which PNSB are known to perform photosynthesis. Fast-growing colonies were isolated, grown anaerobically in 96-well plates containing M22+ broth, and subjected to spectrometry analysis using a microplate reader (700-900 nm scanning; SpectraMax iD3 Multi-Mode). By selecting the fast-growing colonies, we aimed to isolate mutants that were more photosynthetically efficient than the wild-type strain. The plasmids from strains showing spectral characteristics different from the wild-type were purified and submitted for Sanger sequencing. In addition to these random mutants, an amino acid substitution (α-Tyr44Phe and Tyr45Leu) leading to a blue shift of the LH2 absorption peak from 850 to 820 nm described previously (24) was reproduced as a control. The primers used to generate this mutant are listed in Table S1.

### Protein extraction

LH2 variants of interest identified after selection were purified for spectral characterisation as previously described (30). In brief, cells of Rb.X1 at the late-exponential phase of growth and expressing LH2 recombinant variants were lysed by sonication, the cell debris was removed by centrifugation and the supernatant collected. To purify the intracytoplasmic membrane (ICM) containing the photosynthetic proteins, the supernatant was ultracentrifuged twice at 100,000 g for 1 h at 4 °C and water-soluble proteins adhering to the membrane were removed by resuspending the pellet in buffer containing 300 mM NaCl and repeating the ultracentrifugation step. The membrane pellet obtained was then solubilized in solubilization buffer containing Lauryldimethylamine oxide (LDAO) under gentle agitation for 2 h at 4 °C. All subsequent procedures were conducted on ice, in the dark. This solution was ultracentrifuged again and the supernatant containing the proteins, now incorporated into detergent micelles was collected. LH2 was purified from other membrane proteins using a continuous sucrose gradient (0.2-0.6 M). Sucrose solutions were prepared in 20 mM Tris-HCl (pH 8.0) buffer containing 0.2% (v/v) LDAO. The initial membrane protein solution was diluted in the same buffer to adjust the LDAO concentration to 0.2% and layered on the top of the sucrose step gradient. The gradients containing protein samples were then ultracentrifuged at 170,000 g and 4 °C for 16 h, until a protein band, mainly composed by LH2, localized at 0.3 M sucrose could be observed and collected. Next, LH2 preps were further purified by anion exchange chromatography using a HiTrap Q Fast-Flow column (Cytiva). The purity of the final samples containing purified LH2 was estimated based on their absorption at 850 and 250 nm and those with a ratio 850/250 nm > 3 were selected for concentration with an Amicon Ultra-2 Centrifugal Filter Unit (10 kDa MWCO). The concentration of LH2 in solution was calculated using the Beer-Lambert law using 2.9 (mg/ml)^-1^ x cm^-1^ as the extinction coefficient (ε) of LH2 (31).

### Absorption and fluorescence spectroscopy

The spectral characteristics of the LH2 variants obtained were measured from purified proteins preserved in 20 mM tris-HCl (pH 8) buffer with 0.2% LDAO. Absorption spectra were obtained using a GENESYS 30 Visible Spectrophotometer (Thermo Scientific). The emission spectra and the fluorescence decay of the respective peaks were obtained using micro-fluorecence spectroscopy (32). This method uses a confocal microscope coupled to a spectrometer that allows the characterization of the sample’s fluorescence, and single photon avalanche photodiodes (SPADs) to perform time-resolved (lifetime) measurements of emission recombination dynamics.

The fluorescence measurements were performed at room temperature on LH2 samples normalized to a concentration of 2.4 µM in solution, resulting in an average LH2-LH2 distance of 90 nm, thus preventing any intermolecular energy transfers (33). A laser emitting at 472 ± 7 nm, with an 80 MHz repetition rate and pulse width of 60-70 ps was used as the excitation source. The laser spot size was approximately 1.2 μm in diameter, with an excitation power density of approximately 1 μW/μm^2^. The wavelength of 472 nm coincides with the absorption of the spheroidene cars, which in LH2 can transfer the excitation energy directly to B850 through different pathways (34). The fluorescence signal was dispersed using a 0.5 m long spectrometer, with a 600 lines/mm grating blazed at 750 nm, and it was recorded by a silicon charged coupled device (CCD). A SPAD with a measured time response of 340 ps was used as a detector for the lifetime measurements, carried out with a spectral window of 1.5-1.8 nm. The fluorescence lifetime was calculated by fitting the data obtained to a mono-exponential decay in IGOR Pro (WaveMetrics).

### Growth assays

The growth of the Rb.X1 lacking (negative control) or bearing (positive control) the native LH2, mut_860 and mut_820, was monitored in stationary cultures in M22+ broth supplemented with km in 96-well plates at 30 °C. Anaerobic conditions were created within a plastic pouch (Oxoid Compact Plastic Pouch) by an anaerobic gas generator system (Thermo Scientific Oxoid AnaeroGen Compact Sachet) and illumination provided by a VIS-NIR light box (ColorDyne Infrared Waveguide Rod Lamp Mk1 - Blueside Photonics). The experiment was performed at oversaturating light intensities (2000 lux). Optical density measurements at 680 nm (OD680) of each culture were taken with a microplate reader (CLARIOstar® Plus; BMG LabTech) 6 days after inoculation and until the cultures reached the stationary phase. A one-way ANOVA with post-hoc Tukey’s multiple comparisons test was performed to calculate statistically significant differences between the values obtained.

## Results

### Generation LH2 mutants with distinct spectral properties

A library of LH2 mutants of *Rb. sphaeroides* was generated by conducting the random mutagenesis of DNA encoding three target amino acids present in LH2. The library is composed by sequences with single mutations of these amino acids in which all substitutions are possible. The target residues were chosen based on their proximity to the porphyrin ring of the bchlas composing these proteins and the alignment of LH2 homologs identified in different PNSB species (Table S2). In the α-chain, the residues selected were α-Leu20 which makes the closest interaction with the 800-bchla-α through its ester group of the porphyrin E-ring at a distance of 4.9 Å, and α-Leu41 which interacts with 850-bchla-α from a distance of 3.7 Å to its methyl group and 4.1 Å to the alkene in its porphyrin’s B-ring. The residue selected for mutation in the β-chain was β-Thr29 which interacts with the methyl group at the porphyrins’ D-ring of the 800-bchla with a distance of 4.6 Å β-29. Out of them α-Leu20 is conserved in all the species of PNSB being compared, while the other two residues slightly diverge. Notably, β-Thr29 is not conserved in two of the variants encoded by *Rhodopseudomonas palustris* (Table S1).

The expression and successful assembly of the LH2 variants obtained, in addition to the recovery of the native protein in Rb.X1 (221BAC), was confirmed by the presence of absorption peaks at 800 and 850 nm in the mutants grown in 96 well plates. While most of the strains obtained did not display any visible spectral variations, we identified an interesting strain for one of the α-Leu41 mutants displaying a peak red-shifted by 6 nm compared to the B850 absorption features of the native protein. This strain, now called mut_860, was isolated and the plasmid containing the *pucBA* genes coding for this LH2 variant extracted and sequenced. The sequencing results revealed that this red-shifted mutant possesses a substitution Leu41Phe in the α-chain. A known control mutation leading to the expression of an 800-820 nm LH2 variant was also successfully reproduced (24), here referred to as mut_820. The *pucA* gene coding for the α subunit of this variant contained the mutations Tyr44Phe and Tyr45Leu producing a protein similar to the LH3 expressed by *Rbl. acidophilus*. These LH2 spectral variants and the native protein were purified, and their absorption spectra compared side by side (Fig. 2A). Consistent with the shifts observed during selection, the ‘B850’ absorption in the native LH2 peaked at 848 nm, while for mut_820 and mut_860 the maxima of the absorption spectra were measured at 816 and 854 nm, respectively.

**Figure 2.**
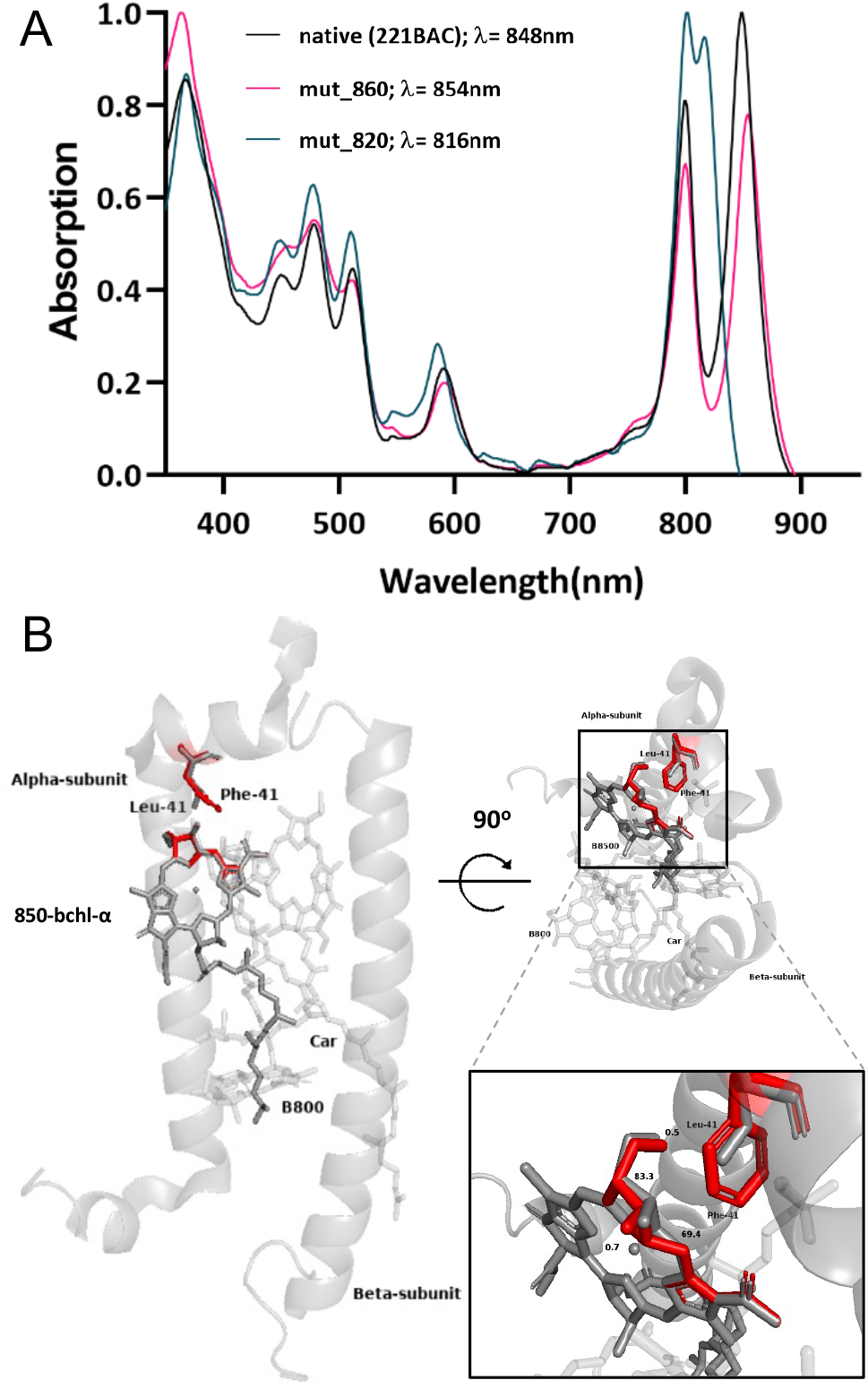
Effect of amino acid substitutions in the absorption properties. **A**. Absorption spectra of LH2 variants. The figure displays the absorption spectra of purified native LH2 and variants obtained by PCR site-directed mutagenesis. Black: native LH2 (221BAC); pink: blue-shifted mutant (mut_820); green: red-shifted mutant (mut_860). Data normalized for maximum absorption = 1.0 with wavelength indicated in the legend for each variant. **B**. Structural prediction of the red-shifted LH2 mutant mut_860 containing the α-Leu41 mutation. The inset zooms in the affected region. The native arrangement of the pigments found between the α and β apoproteins is shown in grey and the changes predicted to occur in this arrangement are shown in red. The structure was generated with PyMOL.

The structural prediction of the red-shifted variant (mut_860) revealed that, compared to the native protein, the mutation present in α-41 affects the interaction of the protein scaffold with the 850-bchla by changing the angle of its porphyrin ring as shown in Fig. 2B. Based on these predictions, this mutation is expected to exert an effect on the porphyrin’s B-ring, causing a bend in this region of the macrocycle towards the second 850-bchla with an angle of approximately 83 Å, displacing the its methyl group by 0.7 Å. Furthermore, this mutation also affects the A-ring of the porphyrin’s macrocycle.

### Fluorescence variations correlate with shifted absorption spectra properties

Next, we investigated how the environments of the photoexcited states of the modified LH2 complexes had been affected by the mutations, by measuring their fluorescence decay rates. We conducted fluorescence lifetime measurements of the different LH2 variants when excited with a laser emitting at 475 nm which corresponds to the spheroidene’s S2 excited state. The fluorescence spectra obtained for the purified native LH2, mut_820, and mut_860 are shown in Fig. 3A. These samples display a fluorescence maximum of 854 nm, 824 nm, and 859 nm, respectively, that corresponds to a Stokes shift of 6 nm, 8 nm, and 5 nm, when compared to the absorption spectra shown in Fig. 2A. A small fluorescence peak is also visible around 800 nm in all samples, corresponding to the fluorescence of the B800 ring of bchlas. The sample presenting the most intense emission is mut_820, while mut_860 and the native LH2 protein (221BAC) present similar but lower (∼ 20%) emission intensities. The variant mut_860 showed slight broadening of the emission spectrum compared to the wild-type. This suggests a greater effect of vibronic mechanisms in the emitting state of this LH2 complex (see discussion section).

**Figure 3.**
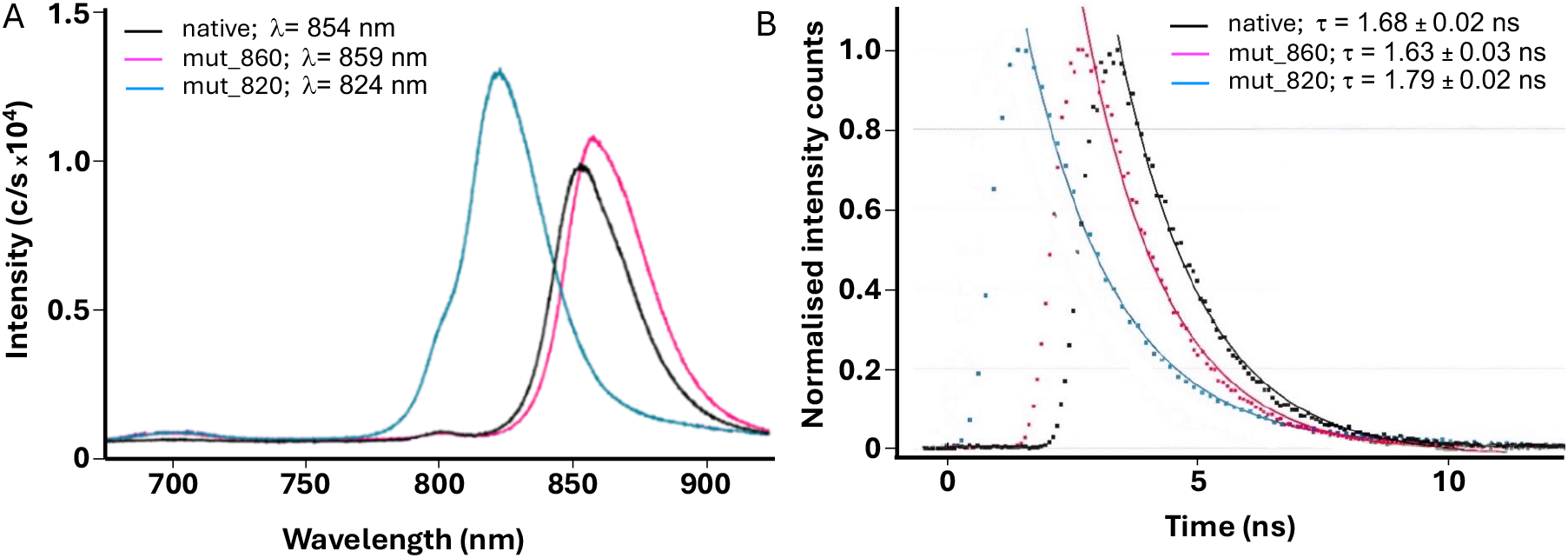
(A) Emission spectra of LH2 variants. The figure displays the fluorescence emission of purified native LH2 and variants obtained through site-directed mutagenesis. Black: native LH2 (221BAC); green: blue-shifted mutant (mut_820); pink: red-shifted mutant (mut_860). Excitation wavelength: 475 nm. Data normalized for an LH2 concentration of = 2.4µM. Wavelengths of emission maxima (/∴) are shown in the legend for each of the variants. **(B) Fluorescence lifetime of LH2 variants**. The figure displays the mono-exponential fits (solid lines) of the time-resolved fluorescence spectroscopy (symbols – experimental data) obtained for the purified LH2. The fluorescence lifetime (ρ), extracted from the fit, is shown in the legend.

We further investigated the fluorescence characteristics of each sample by measuring their emission lifetimes (Fig. 3B). The decay rates _4_were obtained from the fit of the time-resolved fluorescence data collected from each sample to a mono-exponential decay curve. The lifetime measured for the native LH2 of *Rb. sphaeroides* was 1.68 ± 0.03 ns. When compared to the native LH2, the red shifted mutant (mut_680) displayed a lifetime of 1.63 ±0.02 ns, therefore showing an average decrease of 50 ps when compared to the native LH2. The fluorescence lifetime obtained for the native protein compared to the blue shifted mutant (mut_820), instead, displayed lifetime of 1.79 ± 0.02 ns, therefore showing an average increase of 100 ps, showing a. These results indicate a change in the energy transfer dynamics occurring within each LH2 variant in relation to the native protein.

### Absorption spectral variations influence growth phenotypes under high-light intensities

To analyse whether the spectral shifts displayed by the different LH2 variants obtained by directed mutagenesis correlated with phenotypic changes with the potential of affecting the fitness of *Rb. sphaeroides in vivo*, we performed OD680 end-point measurements of cultures of the strains bearing each variant after six days of growth under anaerobic oversaturating light intensities. The results obtained are shown in Fig. 4. The negative control, which was the LH2 deletion mutant Rb.X1 bearing the empty plasmid, reached the lowest cell density (OD680 = 1.204 ± 0.280) as expected.

**Figure 4.**
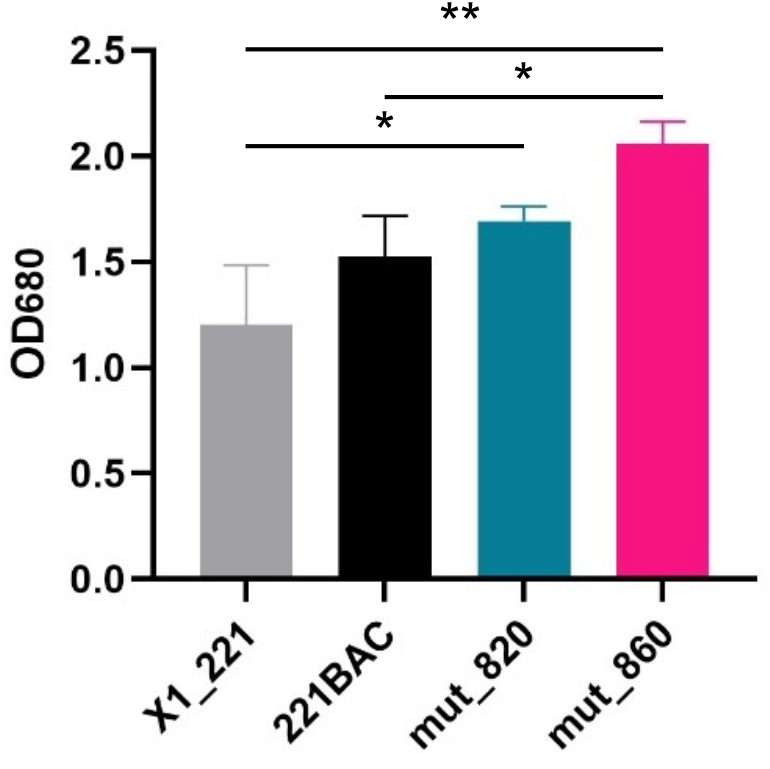
Growth of recombinant *Rb. sphaeroides* expressing LH2 mutants. End-point measurements obtained for *Rb. sphaeroides* strains growing anaerobically under high-light intensity (2000 lux) for six days. X1_221: Rd.X1 transformed with empty pSEVA221 (negative control); 221BAC: Rb.X1 expressing native LH2 (positive control); mut_820: Rb.X1 expressing blue shifted LH2 (mut_820); mut_860: Rb.X1 expressing red shifted LH2. Error bars represent the standard deviation between biological triplicates. (*p<0.05; **p<0.01; ***p<0.001).

Despite a degree of variability in the measurements, there were statistically significant differences observed between the positive control 221BAC expressing the wild-type *pucBA* genes and mut_860, with the latter reaching the highest optical density (2.056± 0.108) followed by mut_820 (1.692 ± 0.069) and the positive control (1.523 ± 0.195). This suggests that the expression of mut_860 can improve the growth of the strains at oversaturating light regimes. This improved growth might be related to an increased resistance to high-light intensity, which is known to be stressful for *Rb. sphaeroides* (35).

## Discussion

The relationship between the complex structure of LHs and their spectral characteristics in PNSB has been widely explored (see for instance (12, 20, 36–39)). In this work, we build on that research to develop a pipeline that could be used for systematic investigations of links between genotype and phenotype of these proteins. We have successfully implemented a workflow for the generation of *Rb. sphaeroides* mutants, expressing different LH2 variants and the selection of mutants of interest for further analysis. We have focused on generating LH2 variants with random mutations in specific amino acids likely to result in spectral changes that has allowed for the selection of fast-growing mutants under photosynthetic conditions. The selected mutants were then screened for the presence of spectral changes based on their absorption spectra and were subsequently characterised in growth assays and by fluorescence spectroscopy.

In our study, the introduction of substitutions similar to the established *Rb. sphaeroides* generated mut_820 (24) produced a B820-like variant with a 32 nm blueshift compared to the native proteins. However, this variant lacks the hydrogen bond formed by α-Tyr41 in LH3 (20, 36), and structural differences like the α-Ser27 interaction with β-bchla (20) may lead to distinct energy transfer dynamics when compared to the native LH3 expressed by *Rbl. acidophilus*.

We have also generated a novel spectral variant, designated mut_860, by substituting α-Leu41 with Phe, resulting in a 6 nm redshift in the absorption wavelength, compared to the native protein. Redshifts in LH2 are rare, and to our knowledge, the 3 nm shift in the LH2 B850 Qy transition linked to the substitution of a serine in position α-27 with a glycine, is the only one reported to date (39). Our LH2 variant mut_860, has the largest redshift reported so far. It has been previously established that B850 redshifts are mostly associated to excitonic coupling among bchlas with some influence of CT state formation (14). The shift observed in mut_860 likely arises from slight porphyrin bending and methyl group repositioning as illustrated in Fig. 2B. Interestingly, α-Phe41 is conserved across several PNSB LH2 types expressed under high-light intensities (Fig. S2), suggesting adaptive relevance. In contrast, in the LH3 of *Rbl. acidophilus*, the blueshift in B820 arises from α-Tyr44Phe and α-Trp45Leu substitutions, leading to altered hydrogen bonding and porphyrin macrocycle distortion (12). These modifications cause an adjustment on the site-energies of the B850 bchlas and the coupling of their excitonic states to CT states (40–45).

The most prominent feature resulting from comparing the emission spectra of the different variants is the substantial difference in the magnitude of the main peak observed in mut_820 in relation to the native LH2. This increase in the emission peak of mut_820 was previously detected in purified LH2 samples derived from *Allochromatium vinosum* (46), and is a consequence of changes in the bchlas site-energies in B820 that reduce the probability of occurrence of non-radiative processes. In this mutant, fluorescence emission seems to become the dominant de-excitation pathway. This result agrees with quantum chemical analyses demonstrating that the coupling between Qy and CT states are significantly reduced in LH3 in comparison to the native LH2 (40), hence providing experimental support for such theoretical predictions. Moreover, mut_860 displayed a slight broadening of the emission spectrum that suggests a stronger influence of vibronic mechanisms in the emitting state of this mutant antenna. This could be due to several reasons including a possible larger coupling to CT states, the opposite of what happens in mut_820.

Important information about the environment surrounding the fluorophores present in the LH2 complexes can be obtained by analysing their fluorescence decay rates. In our variants we observed a clear difference between the fluorescence lifetimes obtained for the Qy transitions of mut_820 and mut_860 in relation to the native LH2. While mut_860 presents a small decrease (from 1.68 ± 0.02 to 1.63 ± 0.03 ns), mut_820 presents a substantial increase in the lifetime (increasing to 1.79 ± 0.02 ns). An illustrative description of the electronic energy gaps and their correlation with the absorption spectrum is provided by Fig. 5. Our data supports a reduction in the energy gap between the Qx and Qy transitions of the B850 bchlas in mut_820 while mut_860 shows a slight increase. These shifts also affect the energy gap between the ground state and Qy.

**Figure 5.**
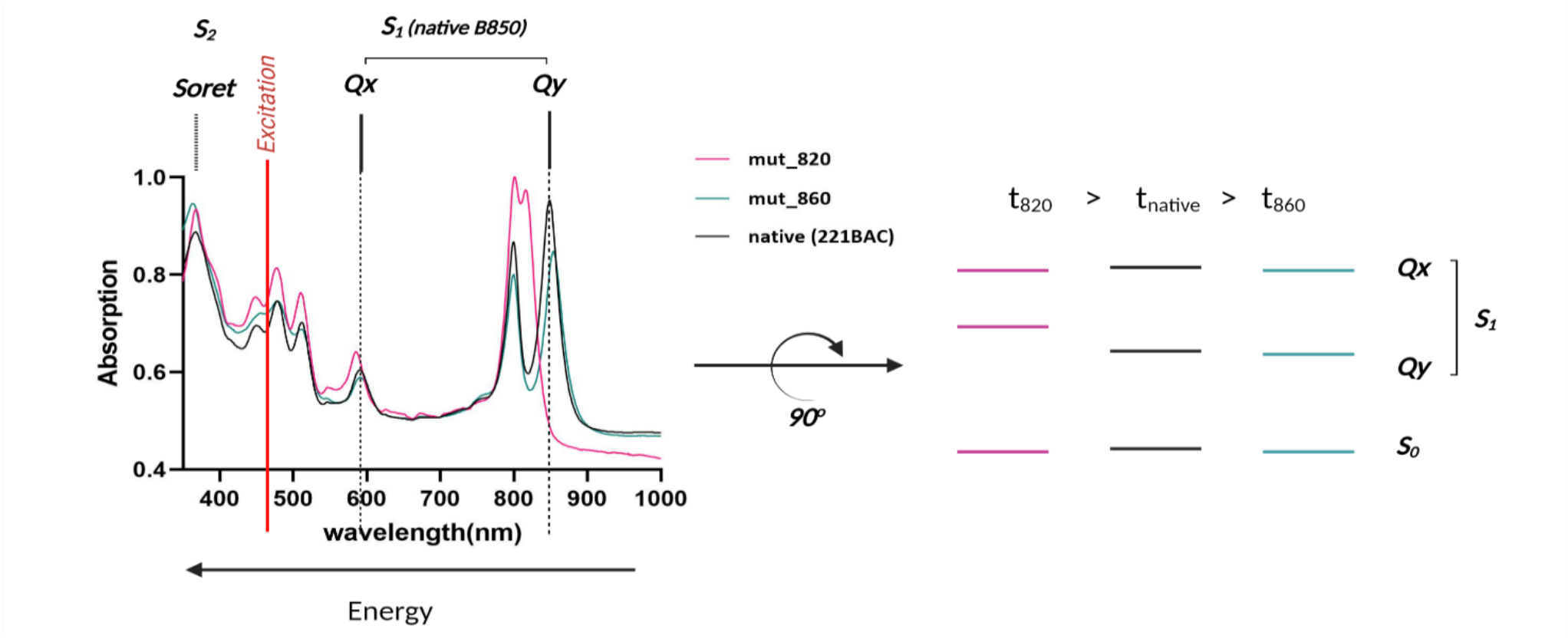
Excited states of LH2 variants. The figure illustrates the correlation between the absorption spectrum and the energy gaps present in the lower energy levels of LH2 (S_1_), Qy and Qx, for the native protein (B850), mut_820 (B820) and mut_860 (B860). The excitation wavelength (475nm) is shown by the red line. Note that for each energy level, only the emitting state (lower excitonic level) is displayed. Pink: mut_820; Black: native protein; Green: mut_860.

Although other interesting processes like intersystem crossing (IC) can play a minor role in the fluorescence lifetime differences observed when comparing each LH2 variant (Gall *et al*., 2011; Kosumi *et al*., 2016; Sipka & Maroti, 2017), we believe that most of the differences observed in our variants can be explained by the phenomenon known as super-radiance. Super-radiance refers to an enhanced radiative rate observed in certain molecular aggregates, in comparison with that of a single monomer, when excited with pulses of wavelength shorter than their separation distance (Jelley *et al*., 1937). For instance, in so-called J-aggregates, that are characterized by a sharp absorption band in the near infrared (NIR) region (also called J-band), relatively short Stokes shifts and fast emission decay rates have been reported (47).

As mentioned previously, the LH2 B850 ring comprises strongly interacting bchlas. The collective nature of its excited states has often been investigated in terms of its super-radiance, which is positively correlated with the degree of excitation delocalization in these aggregates (5, 23, 48, 49). Therefore, it can be hypothesised that most of the decrease in the fluorescence lifetime observed in 860_mut results from an enhancement of the excitation delocalization in these proteins, while the inverse happens in mut_820, when compared to the native LH2. The increase in the radiative lifetime observed in mut_860 would also explain why mutants bearing this variant are able to grow to a higher cell density in high-light regimes. In these mutants, the energy could dissipate faster to the ground state preventing an overload of the RC due to excess radiation and allowing the bacteria to grow efficiently even when exposed to oversaturating light conditions.

The increase of 0.1 ns in the fluorescence lifetime observed in mut_820 agrees with previous findings reporting an increase of around 0.11 ns in the lifetime of the blue-shifted variant of *Rhodospirillum molischianum* in relation to its native LH2 (18). Nevertheless, the evolutionary importance of this spectral shift and the consequent slower rate of fluorescence decay are still unclear. The LH3 complex of *Rbl. acidophilus* is thought to be expressed and increase the fitness of this species in low light conditions due to an enlargement in the energy gap between B820 (LH3) and B875 (LH1-RC) which would diminish the rate of back transfer (18, 50). An *in vivo* experiment carried out under lower light intensities to compare the fitness of each variant in these conditions could provide interesting insights into this open question. However, it is important to note that the artificial conditions created in the laboratory are far from those observed in the strain’s natural environment where many variables can affect its growth.

Further characterisation of the mutants could shed additional light into the dynamics of energy transfer of LH2 proteins. Measurements derived from transient absorption and transient absorption anisotropy would allow the calculation of the energy transfer rates occurring between B800 and B850 and within each ring (18). Additionally, the analysis of circular dichroism spectra would support a better understanding of the interactions occurring between the chromophores, such as the coupling strength between closest neighbours (51) and information about the formation of CT states can be obtained using Stark spectroscopy (52). 2D spectroscopy would provide insights into the exciton delocalization length and coherences occurring between the vibrational modes and the excitonic states of the chromophores (Harel & Engel, 2012; Kim *et al*., 2022). Ultimately, a rigorous analysis of the photoexcited state dynamics in the LH2 variants could be accomplished by obtaining the structure of these proteins using non-invasive techniques such as cryogenic electron microscopy (20).

This study highlights the power of a directed evolution approach to probe and fine-tune LH2 spectroscopic properties. Through targeted mutagenesis and in *vivo* assays employing *Rb. sphaeroides* as model strain, we demonstrate how specific amino acid changes alter photophysical properties of LH2, consequently influencing the growth of PNSB under different light conditions. In addition, our fluorescence decay analysis, supported by structural modelling, provides a starting point to understand energy transfer dynamics in these mutants. This approach can be extended to other systems and can be employed to integrate quantum theoretical predictions with biological experimentation in photosynthesis research. Moreover, our results pave the path for building larger mutant libraries to explore a broader spectrum of spectroscopic and physiological phenotypes. Such libraries would not only deepen our understanding of LH complex adaptation but could also inform the design of synthetic systems for bio-inspired photonic devices.

## Author contributions

E.T.N., JMF, AOC, CAR and JJ designed the study; E.T.N. conducted the experimental work and LS carried out the fluorescence spectroscopy measurements E.T.N and J.J. wrote the paper with contributions from all authors. All authors discussed the results and read, revised, and approved the final version.

## Acknowledgements

We would like to express our gratitude to Prof. Neil Hunter and Dr. David Swainsbury for providing the *pucBA* deletion strain of *Rb. sphaeroides* (Rb.X1) used in this work. This work was supported by the Leverhulme Quantum Biology Doctoral Training Centre, funded by the Leverhulme Trust under Grant no. DS-2017-079. JJ also acknowledges the support received from the Biotechnology and Biological Sciences Research Council (BBSRC) through grants BB/M009769/1 and BB/T011289/1 from the ERA-Cobiotech programme of the EU. LS and AOC acknowledge support from the Engineering and Physical Sciences Research Council (EPSRC) grant EP/V049011/2, and the Gordon and Betty Moore Foundation grant GBMF8820.

## References

1. Madigan, M.T., and D.O. Jung. 2009. An Overview of Purple Bacteria: Systematics, Physiology, and Habitats. In: Hunter CN, F Daldal, MC Thurnauer, JT Beatty, editors. The Purple Phototrophic Bacteria. Dordrecht: Springer Netherlands. pp. 1–15.

2. Li, S., M. Tabatabaei, F. Li, and S.-H. Ho. 2024. A review of green biohydrogen production using anoxygenic photosynthetic bacteria for hydrogen economy: Challenges and opportunities. International Journal of Hydrogen Energy. 54:218–238.

3. Sali, S., and H.R. Mackey. 2021. The application of purple non-sulfur bacteria for microbial mixed culture polyhydroxyalkanoates production. Reviews in Environmental Science and Bio/Technology. 20:959–983.

4. Sauer, K., R.J. Cogdell, S.M. Prince, A. Freer, N.W. Isaacs, and H. Scheer. 1996. Structure-based calculations of the optical spectra of the LH2 bacteriochlorophyll-protein complex from Rhodopseudomonas acidophila. Photochemistry and Photobiology. 64:564–576.

5. Scholes, G.D., and G.R. Fleming. 2000. On the mechanism of light harvesting in photosynthetic purple bacteria: B800 to B850 energy transfer. J. Phys. Chem. B. 104:1854–1868.

6. Harel, E., and G.S. Engel. 2012. Quantum coherence spectroscopy reveals complex dynamics in bacterial light-harvesting complex 2 (LH2). Proceedings of the National Academy of Sciences. 109:706–711.

7. Hu, X., A. Damjanović, T. Ritz, and K. Schulten. 1998. Architecture and mechanism of the light-harvesting apparatus of purple bacteria. Proceedings of the National Academy of Sciences. 95:5935–5941.

8. Hunter, C.N., F. Daldal, M.C. Thurnauer, and J.T. Beatty. 2008. The purple phototrophic Bacteria. Springer Netherlands.

9. Gabrielsen, M., A.T. Gardiner, and R.J. Cogdell. 2009. Peripheral complexes of purple bacteria. In: Hunter CN, F Daldal, MC Thurnauer, JT Beatty, editors. The purple phototrophic bacteria. Dordrecht: Springer Netherlands. pp. 135–153.

10. Cogdell, R.J., A. Gall, and J. Köhler. 2006. The architecture and function of the light-harvesting apparatus of purple bacteria: from single molecules to in vivo membranes. Quarterly Reviews of Biophysics. 39:227–324.

11. Gardiner, A.T., R.J. Cogdell, and S. Takaichi. 1993. The effect of growth conditions on the light-harvesting apparatus in Rhodopseudomonas acidophila. Photosynthesis Research. 38:159–167.

12. McLuskey, K., S.M. Prince, R.J. Cogdell, and N.W. Isaacs. 2001. The crystallographic structure of the B800-820 LH3 light-harvesting complex from the purple bacteria Rhodopseudomonas Acidophila strain 7050. Biochemistry. 40:8783–8789.

13. Fowler, G.J.S., G.D. Sockalingum, B. Robert, and C.N. Hunter. 1994. Blue shifts in bacteriochlorophyll absorbance correlate with changed hydrogen bonding patterns in light-harvesting 2 mutants of Rhodobacter sphaeroides with alterations at α-Tyr-44 and α-Tyr-45. Biochemical Journal. 299:695–700.

14. Fujimoto, K.J., T. Minoda, and T. Yanai. 2021. Spectral tuning mechanism of photosynthetic light-harvesting complex II revealed by ab initio dimer exciton model. J. Phys. Chem. B. 125:10459–10470.

15. Cogdell, R.J., T.D. Howard, N.W. Isaacs, K. McLuskey, and A.T. Gardiner. 2002. Structural factors which control the position of the Qy absorption band of bacteriochlorophyll a in purple bacterial antenna complexes. Photosynthesis Research. 74:135–141.

16. Gardiner, A.T., D.M. Niedzwiedzki, and R.J. Cogdell. 2018. Adaptation of Rhodopseudomonas acidophila strain 7050 to growth at different light intensities: what are the benefits to changing the type of LH2? Faraday Discuss. 207:471–489.

17. Moulisová, V., L. Luer, S. Hoseinkhani, T.H.P. Brotosudarmo, A.M. Collins, G. Lanzani, R.E. Blankenship, and R.J. Cogdell. 2009. Low Light Adaptation: Energy transfer processes in different types of light harvesting complexes from Rhodopseudomonas palustris. Biophysical Journal. 97:3019–3028.

18. Tong, A.L., O.C. Fiebig, M. Nairat, D. Harris, M. Giansily, A. Chenu, J.N. Sturgis, and G.S. Schlau-Cohen. 2020. Comparison of the energy-transfer rates in structural and spectral variants of the B800–850 Complex from purple bacteria. J. Phys. Chem. B. 124:1460–1469.

19. Hördt, A., M.G. López, J.P. Meier-Kolthoff, M. Schleuning, L.-M. Weinhold, B.J. Tindall, S. Gronow, N.C. Kyrpides, T. Woyke, and M. Göker. 2020. Analysis of 1,000+ type-strain genomes substantially improves taxonomic classification of Alphaproteobacteria. Frontiers in Microbiology. Volume 11–2020.

20. Qian, P., T.I. Croll, A. Hitchcock, P.J. Jackson, J.H. Salisbury, P. Castro-Hartmann, K. Sader, D.J.K. Swainsbury, and C.N. Hunter. 2021. Cryo-EM structure of the dimeric Rhodobacter sphaeroides RC-LH1 core complex at 2.9 Å: the structural basis for dimerisation. Biochemical Journal. 478:3923–3937.

21. Gall, A., G.J.S. Fowler, C.N. Hunter, and B. Robert. 1997. Influence of the protein binding site on the absorption properties of the monomeric bacteriochlorophyll in Rhodobacter sphaeroides LH2 complex. Biochemistry. 36:16282–16287.

22. Swainsbury, D.J.K., K.M. Faries, D.M. Niedzwiedzki, E.C. Martin, A.J. Flinders, D.P. Canniffe, G. Shen, D.A. Bryant, C. Kirmaier, D. Holten, and C.N. Hunter. 2019. Engineering of B800 bacteriochlorophyll binding site specificity in the Rhodobacter sphaeroides LH2 antenna. Biochimica et Biophysica Acta (BBA) - Bioenergetics. 1860:209–223.

23. Monshouwer, R., M. Abrahamsson, F. van Mourik, and R. van Grondelle. 1997. Superradiance and exciton delocalization in bacterial photosynthetic light-harvesting systems. J. Phys. Chem. B. 101:7241–7248.

24. Crielaard, W., R.W. Visschers, G.J.S. Fowler, R. van Grondelle, K.J. Hellingwerf, and C.N. Hunter. 1994. Probing the B800 bacteriochlorophyll binding site of the accessory light-harvesting complex from Rhodobacter sphaeroides using site-directed mutants. I. Mutagenesis, effects on binding, function and electrochromic behaviour of its carotenoids. Biochimica et Biophysica Acta (BBA) - Bioenergetics. 1183:473–482.

25. Fowler, G.J.S., S. Hess, T. Pullerits, V. Sundström, and C.N. Hunter. 1997. The Role of βArg-10 in the B800 Bacteriochlorophyll and carotenoid pigment environment within the light-harvesting LH2 complex of Rhodobacter sphaeroides. Biochemistry. 36:11282–11291.

26. Zeng Xiaohua, Choudhary Madhu, and Kaplan Samuel. 2003. A second and unusual pucBA operon of Rhodobacter sphaeroides 2.4.1: Genetics and function of the encoded polypeptides. Journal of Bacteriology. 185:6171–6184.

27. Hunter, C.N., and G. Turner. 1988. Transfer of genes coding for apoproteins of reaction centre and light-harvesting LH1 complexes to Rhodobacter sphaeroides. Microbiology. 134:1471–1480.

28. Silva-Rocha, R., E. Martínez-García, B. Calles, M. Chavarría, A. Arce-Rodríguez, A. de las Heras, A.D. Páez-Espino, G. Durante-Rodríguez, J. Kim, P.I. Nikel, R. Platero, and V. de Lorenzo. 2013. The Standard European Vector Architecture (SEVA): a coherent platform for the analysis and deployment of complex prokaryotic phenotypes. Nucleic Acids Research. 41:D666–D675.

29. Tamura, K., G. Stecher, and S. Kumar. 2021. MEGA11: Molecular evolutionary genetics analysis version 11. Molecular Biology and Evolution. 38:3022–3027.

30. Evans, M.B., A.M. Hawthornthwaite, and R.J. Cogdell. 1990. Isolation and characterisation of the different B800–850 light-harvesting complexes from low- and high-light grown cells of Rhodopseudomonas palustris, strain 2.1.6. Biochimica et Biophysica Acta (BBA) - Bioenergetics. 1016:71–76.

31. Kwa, L.G., D. Wegmann, B. Brügger, F.T. Wieland, G. Wanner, and P. Braun. 2008. Mutation of a single residue, β-glutamate-20, alters protein–lipid interactions of light harvesting complex II. Molecular Microbiology. 67:63–77.

32. Trojak, O.J., S. Gorsky, F. Sgrignuoli, F.A. Pinheiro, J.D. Song, L.D. Negro, and L. Sapienza. 2023. Cavity quantum electrodynamics with quantum dots in aperiodic photonic devices. In: CLEO 2023. San Jose, CA: Optica Publishing Group. p. FF3G.5.

33. Jang, S.J. 2018. Robust and fragile quantum effects in the transfer kinetics of delocalized excitons between B850 units of LH2 complexes. J. Phys. Chem. Lett. 9:6576–6583.

34. Hörvin Billsten, H., J.L. Herek, G. Garcia-Asua, L. Hashøj, T. Polívka, C.N. Hunter, and V. Sundström. 2002. Dynamics of energy transfer from lycopene to bacteriochlorophyll in genetically-modified LH2 complexes of Rhodobacter sphaeroides. Biochemistry. 41:4127–4136.

35. Gürgan, M., H. Koku, I. Eroglu, and M. Yücel. 2020. Microarray analysis of high light intensity stress on hydrogen production metabolism of Rhodobacter capsulatus. International Journal of Hydrogen Energy. 45:3516–3523.

36. Sturgis, J.N., V. Jirsakova, F. Reiss-Husson, R.J. Cogdell, and B. Robert. 1995. Structure and properties of the bacteriochlorophyll binding site in peripheral light-harvesting complexes of purple bacteria. Biochemistry. 34:517–523.

37. Olsen, J.D., J.N. Sturgis, W.H.J. Westerhuis, G.J.S. Fowler, C.N. Hunter, and B. Robert. 1997. Site-directed modification of the ligands to the bacteriochlorophylls of the light-harvesting LH1 and LH2 complexes of Rhodobacter sphaeroides. Biochemistry. 36:12625–12632.

38. Braun, P., A.P. Végh, M. von Jan, B. Strohmann, C.N. Hunter, B. Robert, and H. Scheer. 2003. Identification of intramembrane hydrogen bonding between 131 keto group of bacteriochlorophyll and serine residue α27 in the LH2 light-harvesting complex. Biochimica et Biophysica Acta (BBA) - Bioenergetics. 1607:19–26.

39. Silber, M.V., G. Gabriel, B. Strohmann, A. Garcia-Martin, B. Robert, and P. Braun. 2008. Fine tuning of the spectral properties of LH2 by single amino acid residues. Photosynthesis Research. 96:145–151.

40. Cupellini, L., S. Caprasecca, C.A. Guido, F. Müh, T. Renger, and B. Mennucci. 2018. Coupling to charge transfer states is the key to modulate the optical bands for efficient light Harvesting in purple bacteria. J. Phys. Chem. Lett. 9:6892–6899.

41. Nottoli, M., S. Jurinovich, L. Cupellini, A.T. Gardiner, R. Cogdell, and B. Mennucci. 2018. The role of charge-transfer states in the spectral tuning of antenna complexes of purple bacteria. Photosynthesis Research. 137:215–226.

42. Linnanto, J., A. Freiberg, and J. Korppi-Tommola. 2011. Quantum chemical simulations of excited-state absorption spectra of photosynthetic bacterial reaction center and antenna complexes. J. Phys. Chem. B. 115:5536–5544.

43. De Vico, L., A. Anda, V.Al. Osipov, A.Ø. Madsen, and T. Hansen. 2018. Macrocycle ring deformation as the secondary design principle for light-harvesting complexes. Proceedings of the National Academy of Sciences. 115:E9051–E9057.

44. Anda, A., T. Hansen, and L. De Vico. 2019. Qy and Qx absorption bands for bacteriochlorophyll a molecules from LH2 and LH3. J. Phys. Chem. A. 123:5283–5292.

45. Cardoso Ramos, F., M. Nottoli, L. Cupellini, and B. Mennucci. 2019. The molecular mechanisms of light adaption in light-harvesting complexes of purple bacteria revealed by a multiscale modeling. Chem. Sci. 10:9650–9662.

46. Carey, A.-M., K. Hacking, N. Picken, S. Honkanen, S. Kelly, D.M. Niedzwiedzki, R.E. Blankenship, Y. Shimizu, Z.-Y. Wang-Otomo, and R.J. Cogdell. 2014. Characterisation of the LH2 spectral variants produced by the photosynthetic purple sulphur bacterium Allochromatium vinosum. Biochimica et Biophysica Acta (BBA) - Bioenergetics. 1837:1849–1860.

47. Wang, T., S.F. Yelin, R. Côté, E.E. Eyler, S.M. Farooqi, P.L. Gould, M. Koštrun, D. Tong, and D. Vrinceanu. 2007. Superradiance in ultracold Rydberg gases. Phys. Rev. A. 75:033802.

48. Zhao, Y., T. Meier, W.M. Zhang, V. Chernyak, and S. Mukamel. 1999. Superradiance coherence sizes in single-molecule spectroscopy of LH2 Antenna Complexes. J. Phys. Chem. B. 103:3954–3962.

49. Scholes, G.D., I.R. Gould, R.J. Cogdell, and G.R. Fleming. 1999. Ab initio molecular orbital calculations of electronic couplings in the LH2 bacterial light-harvesting complex of Rps. Acidophila. J. Phys. Chem. B. 103:2543–2553.

50. Deinum, G., S.C.M. Otte, A.T. Gardiner, T.J. Aartsma, R.J. Cogdell, and J. Amesz. 1991. Antenna organization of Rhodopseudomonas acidophila: a study of the excitation migration. Biochimica et Biophysica Acta (BBA) - Bioenergetics. 1060:125–131.

51. Linnanto, J., J.E.I. Korppi-Tommola, and V.M. Helenius. 1999. Electronic states, absorption spectrum and circular dichroism spectrum of the photosynthetic bacterial LH2 antenna of Rhodopseudomonas acidophila as predicted by exciton theory and semiempirical calculations. J. Phys. Chem. B. 103:8739–8750.

52. Somsen, O.J.G., V. Chernyak, R.N. Frese, R. van Grondelle, and S. Mukamel. 1998. Excitonic interactions and Stark spectroscopy of light harvesting systems. J. Phys. Chem. B. 102:8893–8908.

