## Supporting information for "Directed evolution and characterisation of light harvesting complexes with altered energy transfer dynamics in purple non-sulfur bacteria"

### Supporting material

Table S1: List of primers used in this study. Mutation and restriction enzyme recognition sites are shown in bold.

| Nam | Sequence |
|---|---|
| **pucBAC_F** | TTTT**GGATCC**CACGGACTGGCTGGCGTGGG (EcoRI) |
| **pucBAC_R** | TTTT**AATTC**CCCGCAAGATGGGGTCCGGAA (BamHI) |
| **F24** | CGCCAGGGTTTTCCCAGTCACGAC |
| **R24** | AGCGGATAACAATTTCACACAGGA |
| **alpha41_F** | TGCTGACGACCACCACCTGG**NNN**CCCGCCTACTACCAAGG |
| **alpha41_R** | CCAGGTGGTGGTGGAGCAGCGTGGATAACGACG |
| **beta29_F** | ATAAGCAACTCATCCTCGGC**NNN**CACGTCTTCGGTGGCATGGC |
| **beta29_R** | GCCGAGGATGAGTTGCTTATGAACTTCTTCGGCTTCGGCA |
| **alpha20_F** | CCGTCGGCGTTCCGCTGTTC**NNN**AGCGCTGCCGTCATCGC |
| **alpha20_R** | GAACAGCGGAACGCCGACGGTCGGTTTCACCACGAGCCA |
| **mut_820.F** | CCACCACCTGGCTGCCCGCC**TTCCTC**CAAGGCTCGGCTGC |
| **mut_820.R** | GGCGGGCAGCCAGGTGGTGGTCGTCAGCACAGCAGCGTGG |

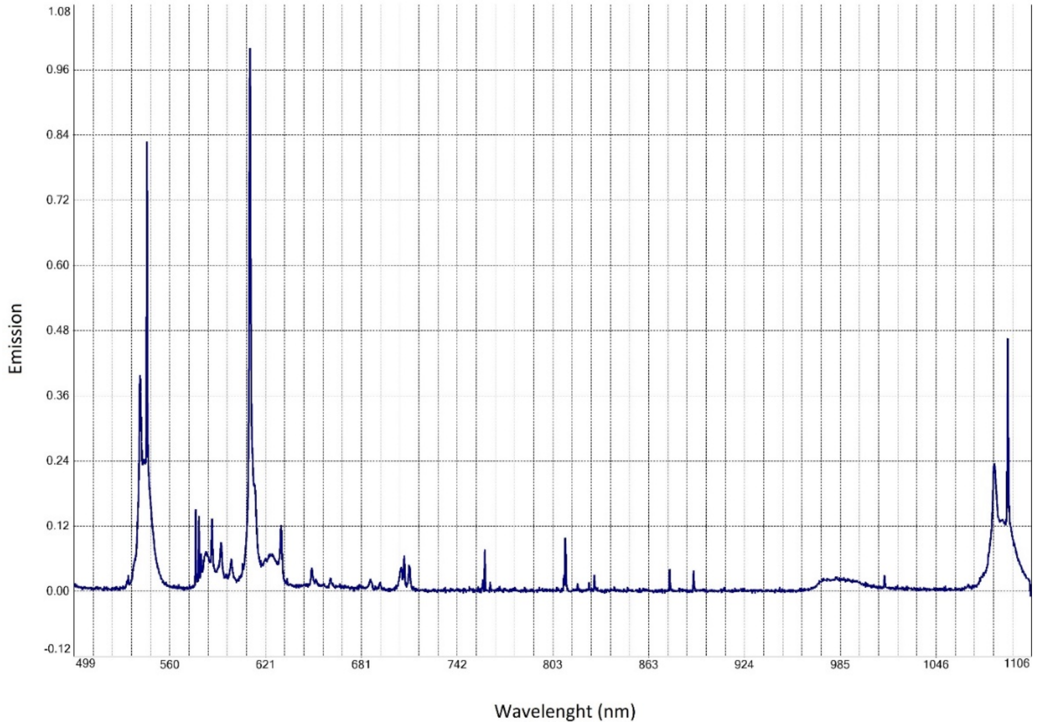


Fig. S1: Anaerobic cabinet's light emission spectrum

a.

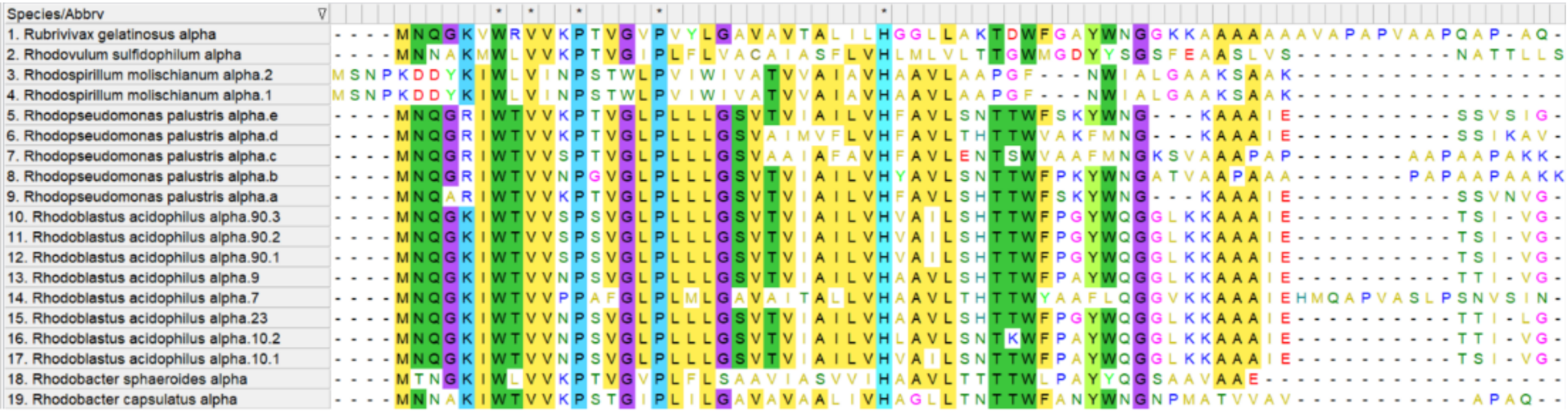


b.

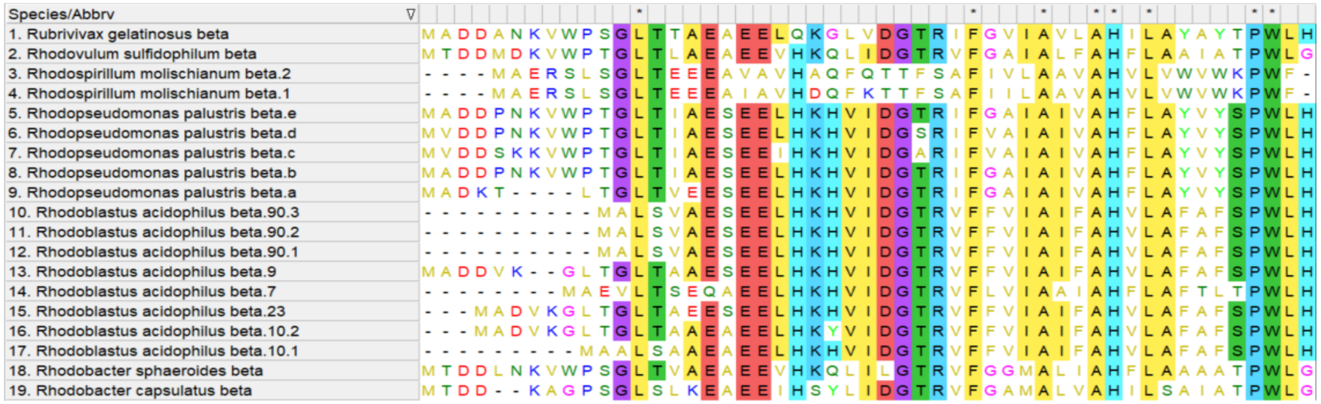


**Fig. S2. Alignment of LH2 subunits**. Sequences of α [a.] and β [b.] subunits correspond to *Rubrivivax gelatinosus* (accession: U67155.1), Rhod*ovulum sulfidophilum* (accession: AAB59006.1 and 7.1), *Rhodospirillum molischianum* (2 variants; accession: CAHP01000001.1), *Rhodopseudomonas palustris* (5 variants; CAA46117.1 to 24.1, CAE26933.1 and CAE26934.1), *Rhodoblastus acidophilus* (8 variants; this study), *Rhodobacter sphaero*ides (accession X68796.1) and *Rhodobacter capsulatus* (accession: PZX26560.1 and 61.1). Amino acids conserved in at least 60% of the species are highlighted. Asterisks mark amino acids with 100% conservation in the species compared.
